# Excel-SBOL Converter: Creating SBOL from Excel Templates and Vice Versa

**DOI:** 10.1101/2022.08.31.505873

**Authors:** Jeanet Mante, Julian Abam, Sai P. Samineni, Isabel M. Pötzsch, Prubhtej Singh, Jacob Beal, Chris J. Myers

## Abstract

Standards support synthetic biology research by enabling the exchange of component information. However, using formal representations, such as the *Synthetic Biology Open Language* (SBOL), typically requires either a thorough understanding of these standards or a suite of tools developed in concurrence with the ontologies. Since these tools may be a barrier for use by many practitioners, the Excel-SBOL Converter was developed to allow easier use of SBOL and integration into existing workflows. The converter consists of two Python libraries: one that converts Excel templates to SBOL, and another that converts SBOL to an Excel workbook. Both libraries can be used either directly or via a SynBioHub plugin. We illustrate the operation of the Excel-SBOL Converter with two case studies: uploading experimental data with the study’s metadata linked to the measurements and downloading the Cello part repository.

**Graphical TOC Entry:** 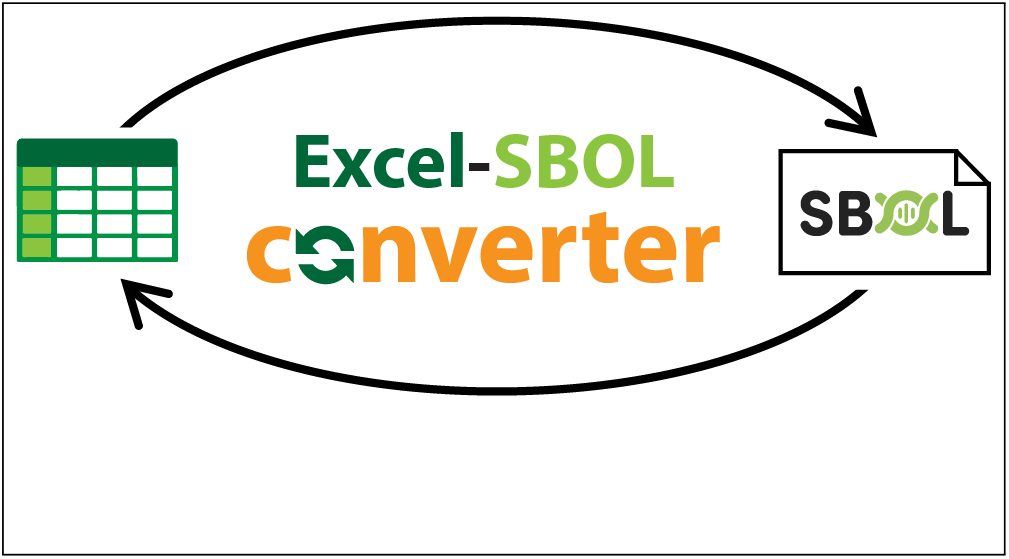

## Introduction

Synthetic biology is bringing together engineers and biologists to design biological circuits for a variety of applications in energy, medicine, and bio-manufacturing.^1^ Associated with this interdisciplinary movement is the need for tools that support reusability and supplement the current understanding of genetic sequences. To satisfy this need, synthetic biology communities across the world have developed tools and ontologies to help describe their unique semantic annotations.^2–15^ Shared representations for data and metadata, grounded in well-defined ontology terms, can help reduce confusion when sharing materials between researchers.^16^ The *Synthetic Biology Open Language* (SBOL)^6^ has been developed to address this challenge. SBOL provides a standardized format for the electronic exchange of information on the structural and functional aspects of biological designs, supporting the use of engineering principles such as abstraction, modularity, and standardization for synthetic biology. Many tools have been created that work with SBOL, ^17–26^ including the SynBioHub repository for storing and sharing designs. ^27^ The original SBOL has been developed further to allow other data types to be represented and more information to be captured. This has lead to SBOL2^28^ and SBOL3. ^29^

Using formal representations such as SBOL, however, typically requires either a thorough understanding of these standards or a suite of tools developed in concurrence with the ontologies.^30^ Unfortunately, this poses a significant barrier for scientists not trained to work with such abstractions. Not using standards makes it difficult to share and reuse parts. ^31–36^ Therefore, the time and effort required to find parts is much greater, and the ability to use the tools to automate design is limited.

One approach to lowering the barrier for use of ontologies was demonstrated by the *Systems Biology for Micro-Organisms* (SysMO) consortium. ^37^ In SysMO, the *MicroArray Gene Expression Markup Language* (Mage-ML) was set up as an XML schema,^38^ and users were expected to submit data to the SysMO Assets Catalogue (called SEEK) in XML format in order to publish work. To allow the use of the Mage-ML language without having to understand XML, the RightField tool was created. ^39^ This tool is an ontology annotation and information management application that can add constrained ontology term selection to Excel spreadsheets. It enables administrators to create templates with controlled vocabularies, such that the scientists utilizing the tool would never actually see the raw RightField, only the more familiar Excel spreadsheet interface. Spreadsheets are a popular interface as many biological workflows already use spreadsheets and *comma separated values* (CSV) files. Furthermore, several popular tools use spreadsheets and CSV files as inputs or outputs, including Addgene (https://www.addgene.org/), and Opentrons (https://opentrons.com/).

Users of SBOL and SynBioHub have also faced a steep learning curve for understanding the underlying ontology: as assessed in,^40^ “For successful use and interpretation of metadata presented in SynBioHub, the semantic annotation process should be biologist-friendly and hide the underlying RDF predicates.” Recently, SynBio2Easy was published as a command line tool to convert Excel spreadsheets of plasmids to SBOL.^41^ The tool was designed to enable several steps of a specific workflow for designing and depositing *Synechocystis* plasmids into a public SynBioHub repository.

Embracing the same spreadsheet-based interface as these prior works, this paper presents the Excel-SBOL Converter, a tool designed to provide a simple way for users to generate and visualize SBOL data without needing a detailed understanding of the underlying ontology and associated technologies. Unlike SynBio2Easy, our converter generalizes beyond plasmids to multiple kinds of SBOL data, as well as allowing customization of the templates. This converter thus provides a simple way for users to manage data by allowing users to both download SBOL into Excel templates and to submit Excel templates for conversion into SBOL. This paper presents the architecture and key engineering decisions for the Excel-SBOL Converter, as well as two case studies that illustrate its operation: uploading experimental data and downloading a repository of parts.

## Results

The Excel-SBOL Converter enables researchers that are more comfortable with Excel spreadsheets to make use of SBOL repositories and tools without having to manipulate or understand the SBOL data standard. The Excel-SBOL Converter is currently implemented as two separate libraries: the Excel-to-SBOL library is used to convert Excel spreadsheets formatted using pre-designed templates into SBOL data, while the SBOL-to-Excel library is used to convert SBOL data into Excel spreadsheets. These two libraries taken together enable data to be converted between Excel spreadsheets and SBOL data as shown in Figure 1.

**Figure 1:**
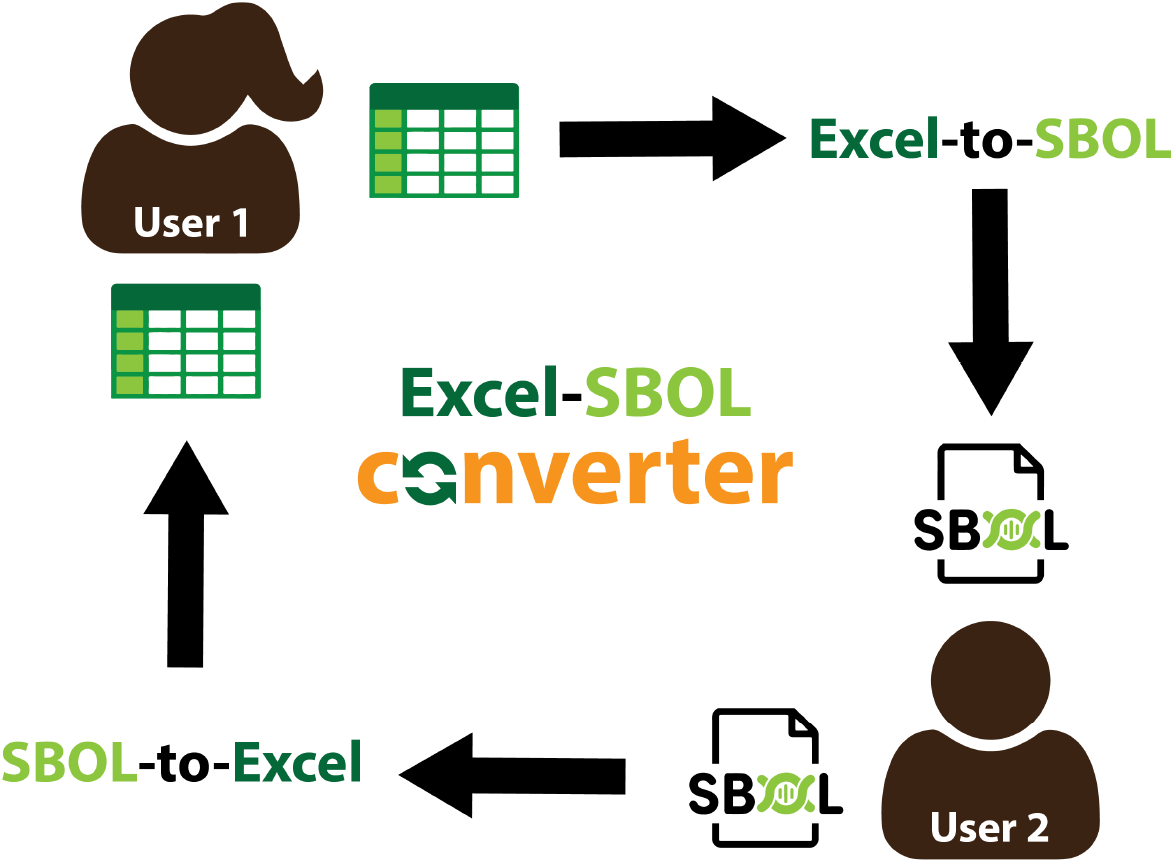
The Excel-SBOL Converter consists of two libraries: 1) Excel-to-SBOL allows the use of Excel templates to create SBOL data, and 2) SBOL-to-Excel allows SBOL data to be converted to Excel spreadsheets. Using the two libraries together allows the creation, uploading, downloading, editing, and re-uploading of SBOL data. Additionally, it allows collaboration between users who prefer spreadsheets and those using software tools that require data to be provided in SBOL format.

### Excel-to-SBOL

The Excel-to-SBOL library converts Excel spreadsheet templates into SBOL data. While initially designed based on fixed spreadsheet templates developed in the Defense Advanced Research Projects Agency (DARPA) Synergistic Discovery and Design (SD2) program (https://sd2e.org/)), the library has since been generalized to allow more flexibility. With this approach, a user needs to have only minimal knowledge of SBOL, while a template designer needs knowledge of both the SBOL data representation and the Excel-to-SBOL template structure.

### Excel Spreadsheet Templates

As an example, let us consider an Excel spreadsheet template that can organize genetic parts into several part collection libraries. This example is composed of one Excel File containing 2 data sheets:

1. A sheet describing the part collection libraries (Figure 2).
2. A sheet describing the genetic parts (Figure 3).

In order to use this template, a user would enter the library names and their descriptions into the collections sheet. The user would then enter each genetic part name, the collection it is a part of, any sequence alterations, part description, data source prefix (PubMed, GenBank etc), Data Source (e.g. the PubMed Id), source organism, whether the part is circular, and the sequence (the final column is automatically generated).

**Figure 2:**
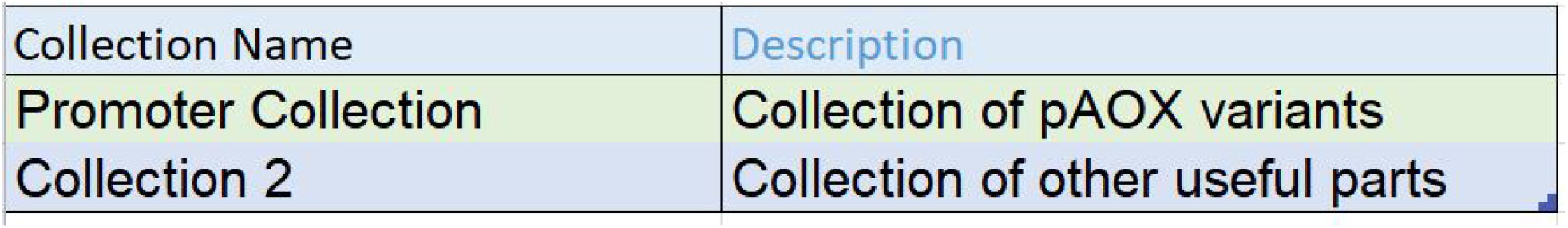
Collections sheet. This is where the users add collections and a description of the collection.

**Figure 3:**
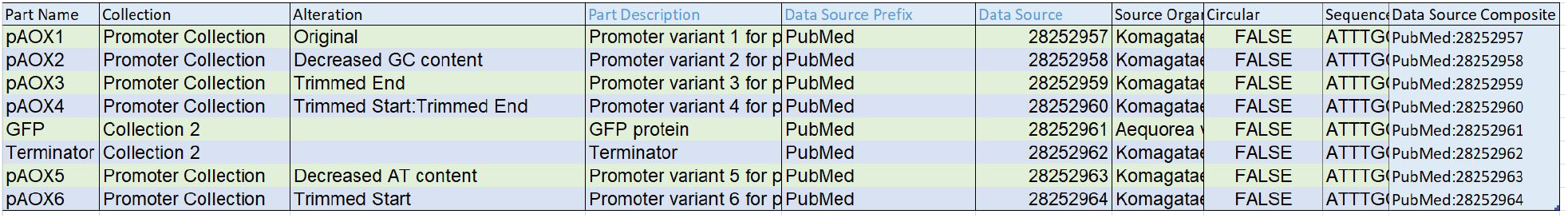
Parts sheet. This is where the users add parts and a description of the parts. Additionally, parts can be added to collections specified in the collections sheet (Figure 2) by adding collection names to the “Collection” column.

### Excel Spreadsheet Template Programming

Excel-to-SBOL is designed to parse a variety of interlinking SBOL object types in a flexible manner, whilst maintaining code simplicity. This is achieved via two main innovations: an initialization sheet, which lists all sheets to be processed, and a column definitions sheet, which provides conversion parameters for each object property.

The initialization sheet is a list of all sheets to be processed (Figure 4). The initialization sheet enables the conversion of multiple sheets. The first column in the initialization sheet is the sheet name, while the rest indicate how it is to be processed. Specifically, these columns indicate whether a conversion should be performed (if True it contains SBOL objects, if False it is ignored except to potentially convert cell values), which row sheet data starts on, whether collection metadata is present (and if so, the number of rows and which columns it can be found in), and whether a collection description is present (and if so, where it can be found). Additionally, further columns may be added to this sheet which are applied to all objects it references. For example, in Figure 4 Molecule Type has been added and filled out for the library sheet. This indicates that all parts on the library sheet have the molecule type DNA.

**Figure 4:**
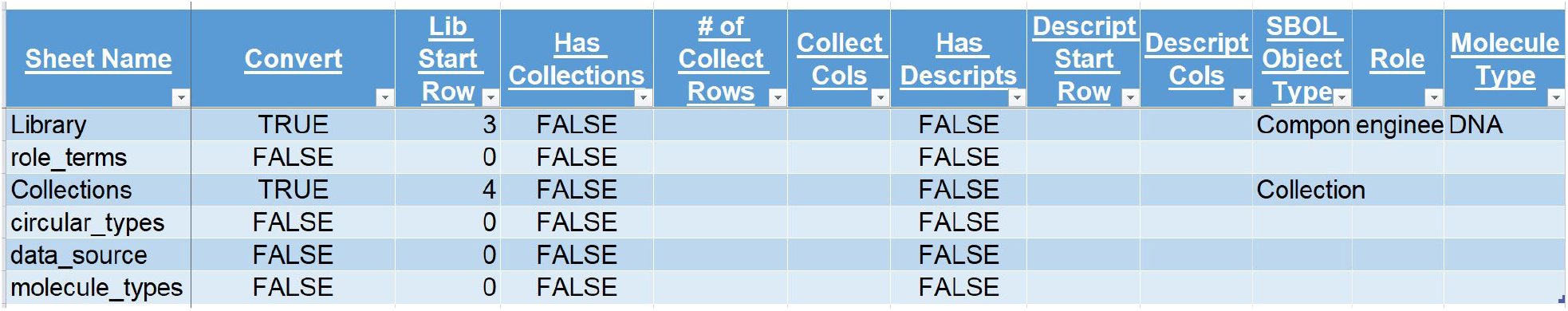
Example initialization sheet. The left most column contains the names of the sheets to be processed. Next the ‘Convert’ column indicates whether the sheet contains SBOL objects (contains SBOL objects if TRUE). The ‘Lib Start Row’ column indicates on which zero-indexed row the information on the sheet starts. The next columns indicate if collection metadata and description are provided or not, and if so, where they can be found. All columns after Descript Cols are columns that contain information that is the same across all items in the sheet.

The column definitions sheet is used for three things: conversion of cell values to a machine readable format, format checking of the cells, and conversion of cell values to SBOL. The column definition sheet for our example is shown in Figure 5. The first two entries identify the sheet and column that conversion is being defined for. The SBOL Term field indicates the property that is found in this location, and how it should be converted into SBOL or a custom data object. This definition begins with a short prefix for the namespace (e.g., “sbol”) followed by an underscore and the property name itself. Most terms will lead to a simple addition or the property, however some (e.g. sbol sequence) call specialized functions that do more than just add a property (e.g. also creates a sequence object). Additional fields indicate the namespace URL (Uniform Resource Locator) to use for the property, the type of the value (e.g., String or URI (Uniform Resource Identifier)), whether the property can have multiple values and the character(s) on which the field is split for conversion into a list of values. Additional value checking can be performed with a “pattern” field that provides a series of regular expressions that the property value is to be checked against, separated by quotations. For example: “ *∧* [*a − zA − Z\s∗*] +$” “*https:\/\/www.ncbi.nlm.nih.gov/nuccore/. ∗* “ is used for checking sequence entries, indicating that entries should either contain only alphabetical characters and spaces or be URLs starting with https://www.ncbi.nlm.nih.gov/nuccore/.

**Figure 5:**
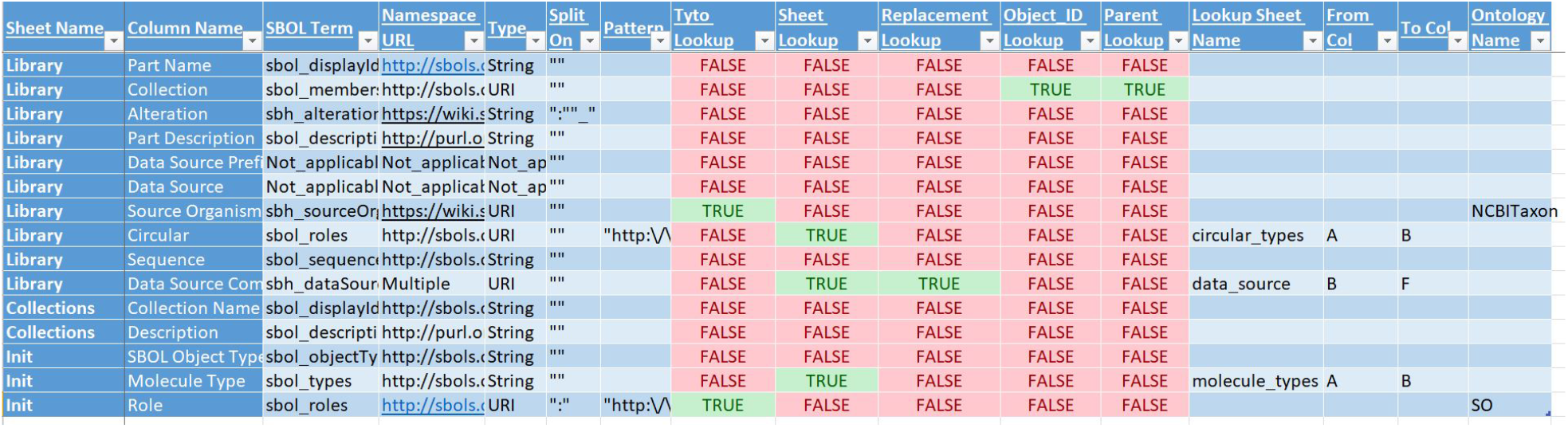
Example column definitions sheet. The Sheet Name and Column Name entries identify the sheet and column within the sheet that is being defined. Next, the SBOL Term indicates the encoding property for the cell. The Namespace URL is the namespace to be used with the encoding property, and the Type column indicates the type of data expected (i.e. URI or String). The Split On column is used to split a single column into multiple values for a property. The Pattern column contains regular expressions to check the cell values after they have undergone conversion. The following columns contain conversion information to go from human to machine-readable values. Tyto Lookup uses the Tyto ^42^ library to perform lookups according to the “Ontology Name” Column. The library performs lookups according to the “Ontology Name” Column. Sheet lookup takes the cell value and converts it to another value like a lookup dictionary. The dictionary is created based on the “Lookup Sheet Name”, “From Col”, and “To Col”. Replacement Lookup is a special case of sheet lookup. In this case, the cell value is expected to be prefix:value. Object ID Lookup converts the cell value to an SBOL Object URI. The “Parent Lookup” column indicates the directionality of the object reference.

The remaining fields are used to define mechanisms that can be used for conversion from human readable to machine readable forms. The first option is to use the Tyto software to convert a string to an ontology term. ^42^ In this case, the ontology to look the terms up in must also be given (“Ontology Name”). For example, in Figure 4, Role (on the Init Sheet) is converted using the *sequence Ontology* (SO).^43^

The second option is to lookup the ontology term using another sheet within the spreadsheet as a dictionary (see for example Figure 6). When this option is selected, the user must also specify the sheet to use for the lookup (“Lookup Sheet Name”), the column in which the human readable value is found (“From Col”), and the column in which the machine readable ontology term is found (“To Col”). For example, following the dictionary shown in Figure 6), TRUE in the Circular column is converted to http://purl.obolibrary.org/obo/SO_0000988 (the Sequence Ontology Role for circular) and false means no additional role is added.

**Figure 6:**
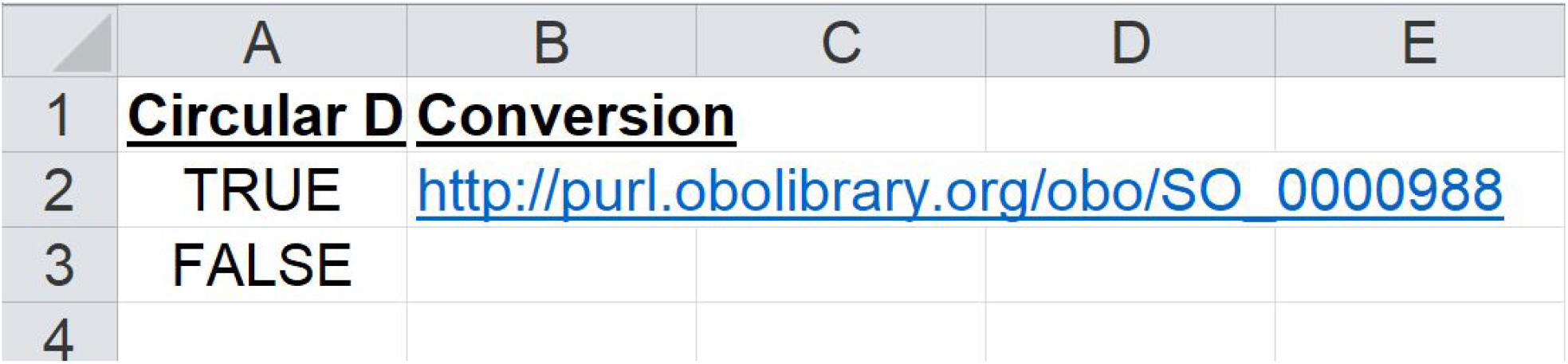
Lookup Sheet (circular types). This spreadsheet is used in a Lookup replacement and converts human readable terms to machine readable ones. In this case the value TRUE in the circular column leads to the Sequence Ontology term for circular whilst FALSE leads to no addtion.

Replacement Lookup is a special case of sheet lookup. In this case, the cell value is expected to be prefix:value, e.g. PMID:24295448, representing the data source is a PubMed id with the value 24295448. Here the lookup works by using the prefix as the key and pulls a value from the “To Col” and inserts the cell suffix in the place of {REPLACE HERE} (Figure 7). So for the case PMID:24295448, the PMID key gives the string https://pubmed.ncbi.nlm.nih.gov/{REPLACE_HERE}, which is changed to https://pubmed.ncbi.nlm.nih.gov/24295448. This kind of lookup is useful in the case where several different kinds of information are put in one column, e.g., a data source might be GenBank, PubMed, AddGene, etc. Additionally, it allows the formatting of a URL rather than being a direct remapping in the way a simple sheet look up is.The final kind of lookup is Object ID lookup. The Object ID Lookup means that the cell value is converted to a URI. For example, a part called GFP promoter can be referenced using that name, but this reference can then be converted to the URI http://www.examples.org/gfp_promoter using this lookup. This type of lookup only works for SBOL objects. If the parent lookup column is also true, then it indicates that the current object is added as a property to the referenced object rather than vice versa. For example, in a row for a component definition called pAOX1 (Figure 3), there may be a collection column with the collection name: Promoter Collection. If the Parent Lookup is true, then rather than adding Promoter Collection to pAOX1 using the sbol member attribute, the pAOX1 URI is added to the Promoter Collection object using the sbol member attribute.

**Figure 7:**
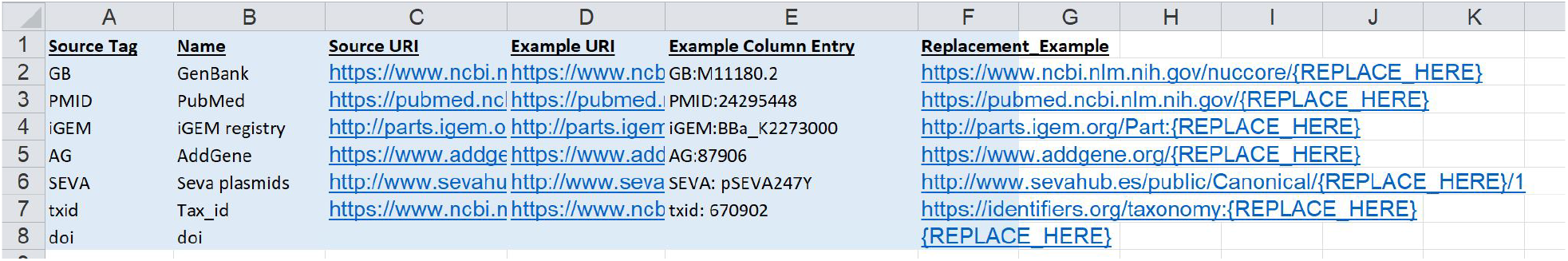
Replacement Lookup Sheet. This is an example of a replacement lookup sheet referenced by the column definitions sheet. In this case the prefix found in column B is used to identify the correct URL in column and the suffix is placed in the URL at the location indicated by {REPLACE_HERE} So for the case PMID:24295448, the PMID key gives the string https://pubmed.ncbi.nlm.nih.gov/{REPLACE_HERE}, which is changed to https://pubmed.ncbi.nlm.nih.gov/24295448.

### Excel-to-SBOL Conversion Algorithm

The initialization and column definition sheets are used by the Excel-to-SBOL conversion algorithm shown in Algorithm 1. The full code for the Excel-to-SBOL converter can be found at https://github.com/SynBioDex/Excel-to-SBOL. This algorithm proceeds as follows. First, it reads in the initialization, column definition, and data sheets. It then adds any additional properties specified on the Init sheet to the column definitions sheet as well as the appropriate data sheet. Next, it creates an SBOL Document, and it begins parsing all the sheets that have been marked in the initialization sheet as containing SBOL content. To do this, it identifies the displayId and SBOL object type columns, and it creates an SBOL object of the specified type for each entry found with a URI constructed from the displayId found. Finally, it parses all the other columns in each sheet as specified in the column definitions sheet. First, it splits the entries found using the Split On field. Next, it converts the value found, if needed, using the lookup method specified in the column definition sheet for that entry. Then, it checks the converted cell value against the regular expression provided in the Pattern field. Finally, it adds the property and value to the corresponding SBOL object using the specified SBOL Term and Namespace URL.

#### Algorithm 1: Excel-to-SBOL

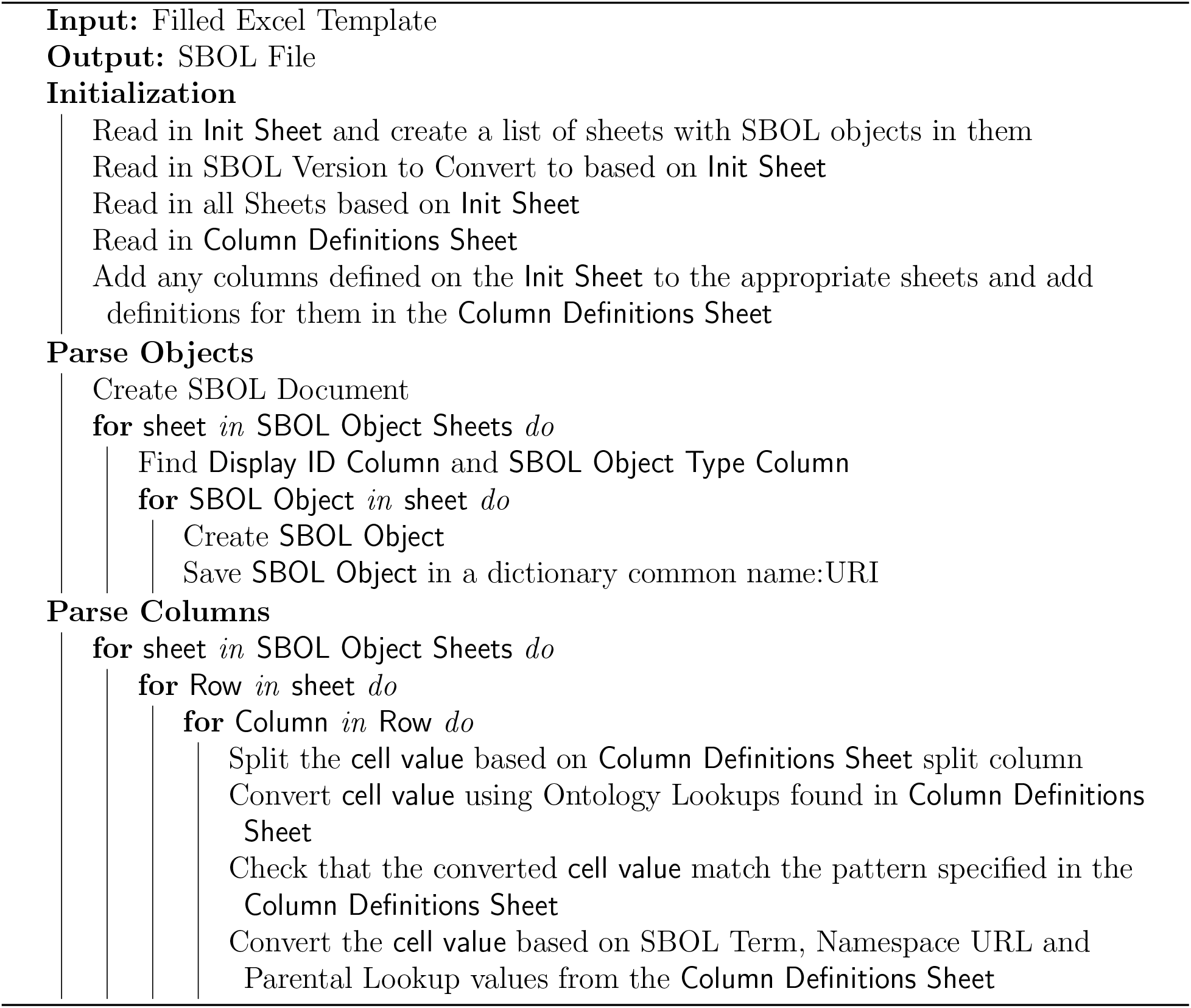

### SBOL-to-Excel

The SBOL-to-Excel converter performs the opposite task of taking SBOL data and converting it into an Excel spreadsheet. This converter reads an SBOL file in any standard RDF format (e.g., XML, JSON-LD, Turdle, N-triples), and for every SBOL object found, it creates a row in a spreadsheet that has a column for each property of the object. In this case, the RDFlib python library is used rather than the more specialized SBOL libraries in order to allow the conversion of different SBOL object types in any version of SBOL whilst maintaining code simplicity.

An example Excel spreadsheet generated by the SBOL-to-Excel converter from the SBOL generated for the library example presented earlier is shown in Figure 8. This spreadsheet looks similar to the templates described earlier, but has a few key differences. First, all column names are based on the property name rather than human readable column names. Second, no Init or Column Definitions sheet is present in the output. Finally, many of the properties are full URLs rather than human readable names. Future work will address these differences to allow full round-tripping of data from Excel to SBOL and back again.

**Figure 8:**
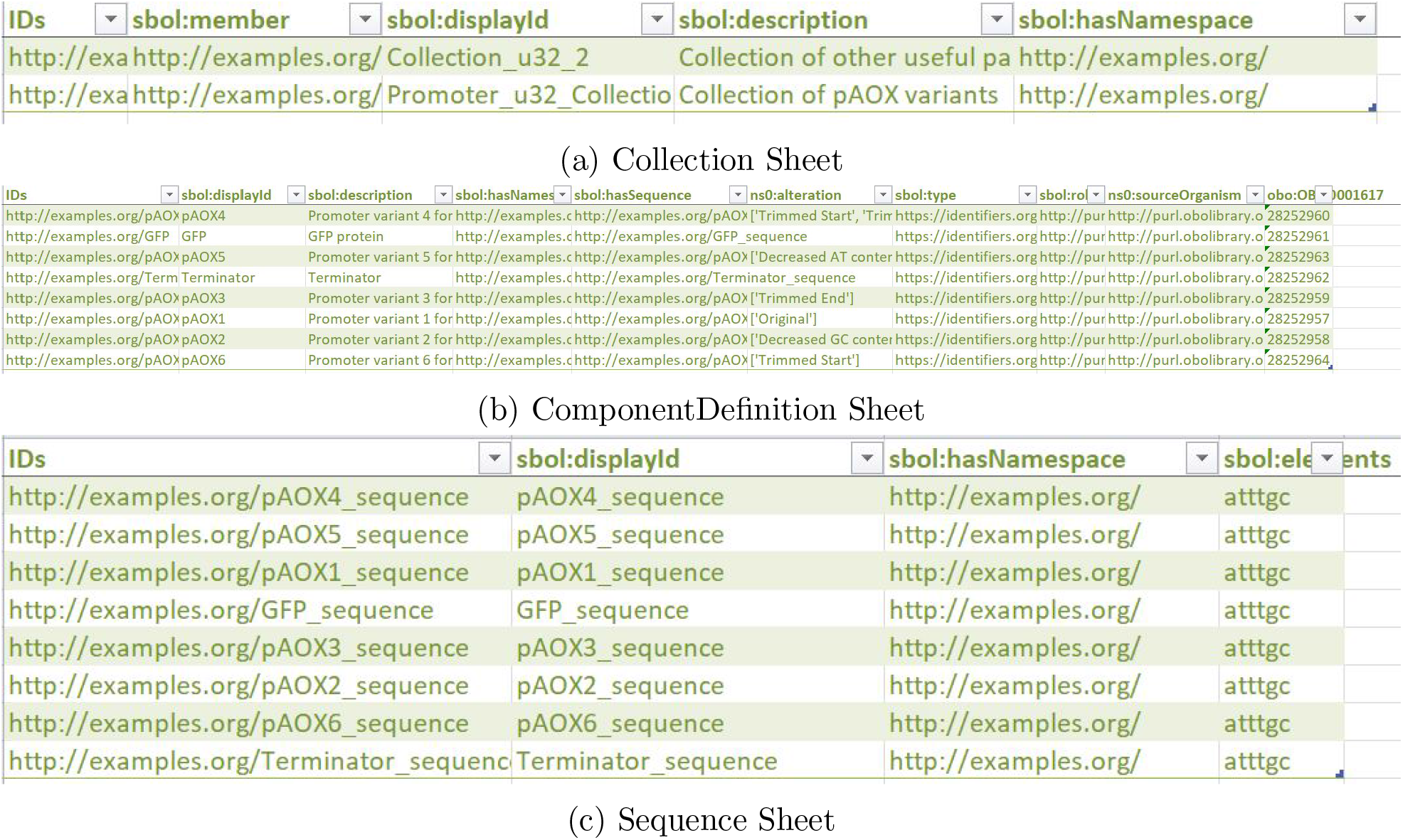
Spreadsheet output by the SBOL-to-Excel Converter. This is an example output for the SBOL example shown in the Excel-to-SBOL section. Note that each SBOL object type is split into separate sheets.

The algorithm used by the SBOL-to-Excel Converter is shown in Algorithm 2. The full code for the SBOL-to-Excel Converter can be found at https://github.com/SynBioDex/SBOL-to-Excel.

The algorithm first loads SBOL data into a dictionary with a dictionary of predicates containing a list of objects for each subject. This is then converted to a pandas DataFrame. Next, the predicate prefixes are converted to more human readable forms (e.g. “http://purl.org/dc/terms“ to ‘dcterms’). Then the columns are reordered according to a hardcoded priority system and any columns in a hard-coded removal list are removed. Finally, each SBOL object type is output to its own sheet.

#### Algorithm 2: SBOL to Excel

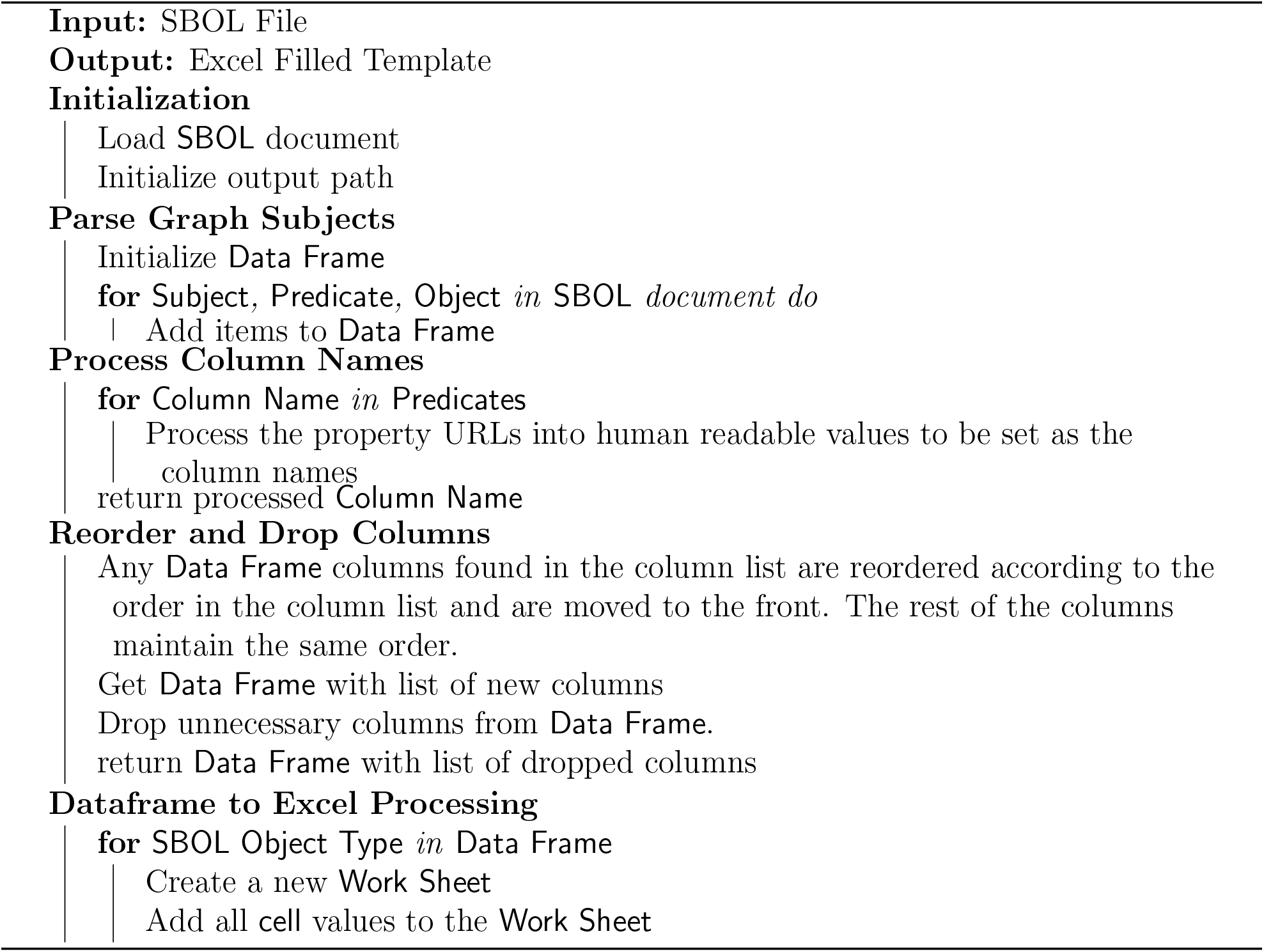

### SynBioHub Plugins

SynBioHub plugins for the Excel-SBOL Converter were developed to enable the converter to be integrated into a data processing workflow using the SynBioHub repository (see Figure 9). SynBioHub plugins are a way to create modular extensions to the capabilities of the SynBioHub repository.^44^ The plugins developed as part of this project allow the integration of SynBioHub into synthetic biology workflows with spreadsheets. A submit plugin was developed for the Excel-to-SBOL converter. This plugin takes in Excel spreadsheet templates, converts them to SBOL, and uploads the converted SBOL to SynBioHub. Similarly, a download plugin was developed using the SBOL-to-Excel library that allows a user to download SBOL stored in SynBioHub in the form of an Excel spreadsheet.

**Figure 9:**
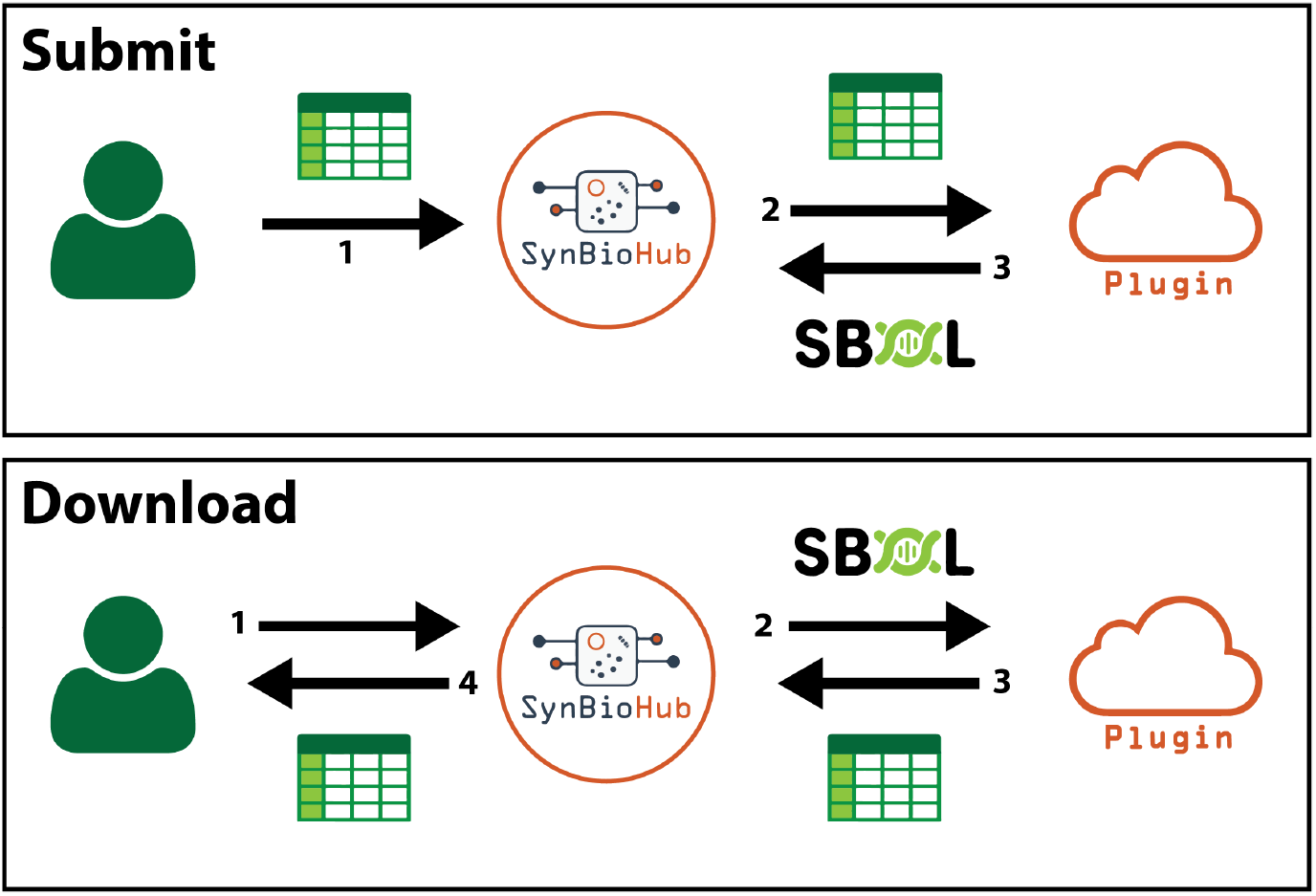
Integration of the Excel-SBOL Converter with SynBioHub via plugins. **Submit**: When a user uploads an Excel spreadsheet template to the submit endpoint, SynBioHub sends it to the Excel-to-SBOL plugin, which returns SBOL to be deposited in SynBioHub. **Download**: If a user requests an Excel spreadsheet template to be downloaded from SynBioHub, SynBioHub sends the appropriate SBOL to the SBOL-to-Excel plugin. The plugin returns an Excel spreadsheet to SynBioHub, which is then returned to the user.

## Case Studies

We now illustrate the operation of the Excel-SBOL Converter with two case studies, one focused on the upload of experimental data, the other focused on downloading a part library.

### Experimental Data Sheets

Capturing experimental data with Excel-to-SBOL allows researchers to more easily associate measurement metadata with genetic part information. The experimental metadata may include descriptions of experimental samples, study conditions, assay types, and equipment. All this information can be captured in the SBOL data standard. To process experimental data, an Excel spreadsheet template was created that includes sheets for Studies (i.e., experimental replicates), Assays (i.e., experiments), Sample Designs (i.e., the media, strain, and vector used), Samples (i.e., an individually measured instance of a sample design), and Measurements (i.e., the data collected).

In the example, shown in Figure 10, each experimental study is conducted over a period of twenty-five hours with measurements taken every 15 minutes. This experimental study is repeated two times, with the study composed of two assays and each assay comprising of two samples with two sample designs, for a total of four samples. Each of these four samples then has two signals (i.e., *optical density* (OD), *green fluorescent protein* (GFP)) measured for 100 time points, resulting in a total of 800 measurements.

**Figure 10:**
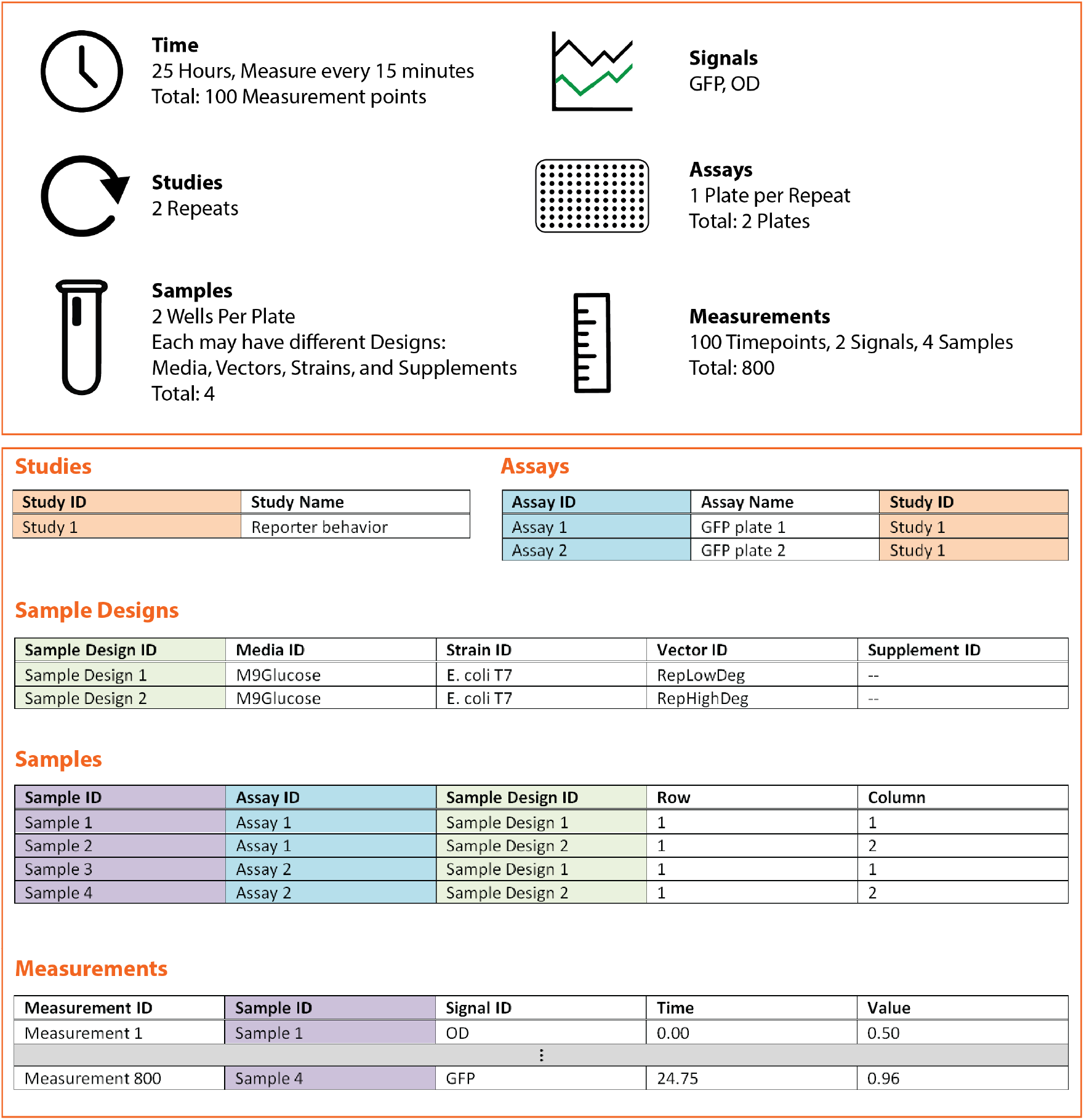
Excel-to-SBOL example for a collection of experiments. **Top**: Diagram showing the design of an example experiment comprising a study with measurements every 15 minutes over a period of twenty-five hours with two repeats. For each repeat, there is one plate per repeat with one assay per plate. Each plate has two samples per assay for a total of four samples at two different sample design setups. Measuring each sample at each of the 100 time points results in 800 data points. **Bottom**: Given this experimental setup, five different sheets are used to describe and connect information for Studies, Assays, Sample Designs, Samples, and Measurements. Note the way the sheets are linked via IDs. For example, assays identify the study they are part of using the “Study ID” column and similarly, samples to assay with “Assay ID”, and measurements to sample with “Sample ID”.

The information in this spreadsheet can be uploaded for storage in the SynBioHub repository^45^ after processing by the Excel-to-SBOL converter. The Excel-to-SBOL converter converts each study into an SBOL Collection object, each assay into an SBOL Experiment object, each sample design into an SBOL Module Definition object, and each sample into an SBOL ExperimentalData object. However, Measurements are not converted to SBOL and are not uploaded to SynBioHub. In the future, we plan to upload the measurement data to the Flapjack repository^46^ (a repository for the storage of experimental measurement data).

### Cello

The Cello library^47^ was chosen as a case study for the SBOL-to-Excel Converter due the variety of SBOL objects that it contains, including objects with a diverse set of custom annotations. In order to achieve compliance with these varied SBOL types, it is key that the SBOL-to-Excel Converter has a way of dynamically manipulating these types. With the XML parsing capabilities offered by RDFLib, the SBOL-to-Excel Converter is able to successfully process and organize all of the data into a 15 sheets in an Excel document. The conversion to Excel facilitates the analysis of the SBOL document for a user that prefers spreadsheets. The results of the Cello conversion can be seen in Figure 11.

**Figure 11:**
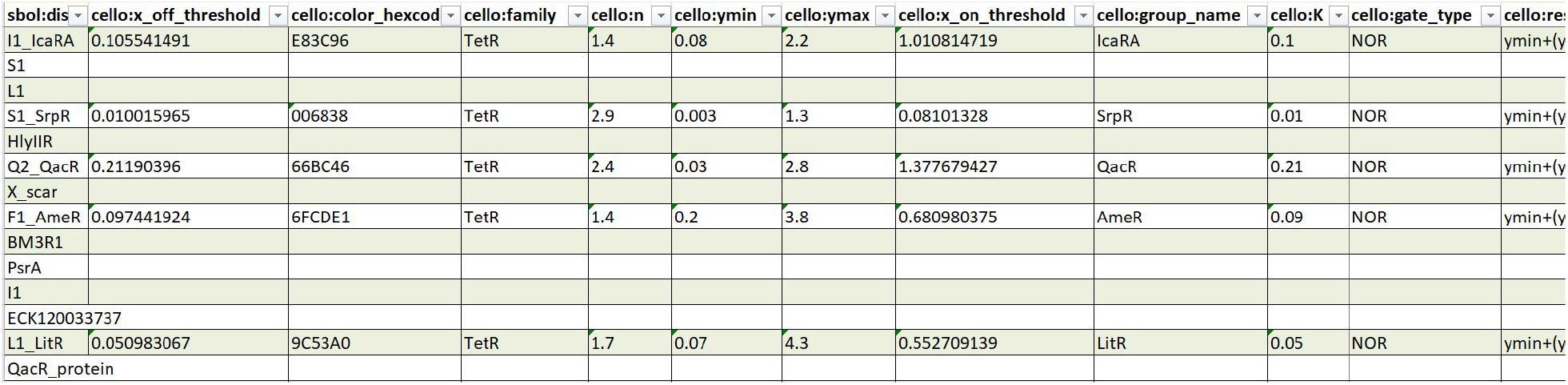
Example of a part of a sheet (ComponentDefinition) output by the SBOL-to-Excel Converter from the conversion of the Cello part Library. ^47^ In total 15 sheets are output for: Range, Collection, Interaction, ComponentDefinition, ModuleDefinition, Participation, FunctionalComponent, Sequence, Attachment, SequenceAnnotation, Component, Activity, Agent, Association, and Usage. Note the human readable column names.

## Discussion

We have presented two Python converter libraries: Excel-to-SBOL and SBOL-to-Excel. The development of these libraries simplifies the incorporation of SBOL into existing synthetic biology workflows that make use of spreadsheets for data storage and exchange. The Excel-to-SBOL converter allows multiple sheets with different SBOL object types to be processed. The converter can be used to convert spreadsheet columns into any RDF properties, and can output either SBOL Version 2 or 3 documents, as well as validating information using user supplied regular expressions and converting user-friendly names into ontology terms. Its complement, the SBOL-to-Excel Converter, can process any RDF file and turn it into a spreadsheet. This allows both SBOL2 and SBOL3 documents to be converted to spreadsheets. The conversion makes the values more human readable, and splits different SBOL object types across sheets.

While the Excel-SBOL Converter is already useful for many applications, there are several improvements planned for the future. For the Excel-to-SBOL Converter, real-time sheet checking using Excel plugins would enable errors to be detected and fixed more efficiently. Second, more standard Excel spreadsheet templates with example data should be created to provide a starting point for both users and template creators. For the SBOL-to-Excel Converter, the next step is to record the conversion process into a column definition sheet in order to support round-trip conversion back to SBOL.

Finally, the case studies presented here both indicate that more thought needs to be given to development of core standards regarding the information to be collected and shared about experiments and parts. There is currently no consensus regarding what information is desirable to share or how to organize it. Excel templates can be made available using the Excel-SBOL Converter, however, it may be possible to help in developing a *de facto* standard for information to be collected, and such a minimum information standard should be further considered.

## Methods

Below is a more in depth description of the methods used by the Excel-to-SBOL and SBOL-to-Excel Converters, as well as the methods used in the case studies. Note that all code was written in Python.

### Excel-to-SBOL

This converter relies on a set of Python modules to function. The modules used and their functionality in the converter is explained below.

- **Openpyxl** is a Python library to read and write Excel files. This library is used together with Pandas to read in the data from the Excel workbook and store it as a data frame.
- **pandas** is a data analysis library. It is used for the reading in of the Excel workbook and the further storage and manipulation of the data.
- **pySBOL2 and pySBOL3** are libraries for the reading, writing, and manipulation of SBOL data. These libraries are used to create the SBOL objects required and write them out to an SBOL document.
- **Tyto** is a tool to make the semantic web more accessible. It is used to convert cell values to ontology terms.

### SBOL-to-Excel

SBOL-to-Excel relies on a set of libraries to ensure the smooth processing of a given SBOL document. These libraries facilitate advanced processing procedures, and ensure that the data is properly output to Excel for the user to analyze.

- **RDFLib** is a python package that enables the user to work with the Resource Description Framework (RDF). RDF serves as an important proponent to this project. Its functionality includes the ability to parse through RDF triples within an Excel file. With access to the subject, object, and predicate triples, the library made it possible to extract these values, and convert them into a form that can be easily interpreted in order to eventually be output to Excel.
- **Openpyxl** is a Python library to read and write Excel files. The Openpyxl library is crucial to the Converter, as its functionality allows the writing of SBOL data into the Excel spreadsheet. It is then also used to format the spreadsheet (including creating the hyperlinks).
- **Pandas** is a python library that provides data structures with capabilities for the accessing and manipulating of data that it holds, and does so with the information processed from the SBOL collection. This occurs specifically when the data is being read in. The subjects, objects, and predicates are set into the pandas data frame in order to be processed by various modules, before being output into Excel.
- **Validators** is used to check items expected to be URLs are valid URLS.
- **Pytest** makes it easy to write small tests, as well as, scales to support testing for large applications. This library enabled modular testing, ensuring that the correct forms of data were being passed through the converter at the appropriate points.

## Supporting information

Excel to SBOL file

SBOL to Excel example

Flapjack Case study file

Cello case study example

## Acknowledgement

JA, JM, and CM are supported by the National Science Foundation under Grant No. 1939892. JM is additionally supported by a Dean’s Graduate Assistantship at the University of Colorado Boulder. JA is additionally supported by a CU Boulder Discovery Learning Apprenticeship. IP was supported by the Google Summer of Code and the Bischöfliche Studienförderung Cusanuswerk. PS was supported by the SBOL Industrial Consortium. JB and CM are partially supported by Air Force Research Laboratory (AFRL) contracts FA8750-17-C-0184 and FA8750-17-C-0229. SynBioHub and the Excel-SBOL Converter plugins are run on a Microsoft Azure Server provided by Microsoft Research. This document does not contain technology or technical data controlled under either U.S. International Traffic in Arms Regulation or U.S. Export Administration Regulations. Any opinions, findings, and conclusions or recommendations expressed in this material are those of the author(s) and do not necessarily reflect the views of the funding agencies. All authors contributed to the writing of this manuscript.

## Author Contributions

All authors contributed to the writing of this manuscript. JM worked on Excel-to-SBOL, supervising IP in the initial design and then taking over. JA worked on SBOL-to-Excel with the help of JM. SS created the experimental data case study. PS is working on Tyto integration into Excel. CM supervised the project. JB contributed guidance and the initial DARPA spreadsheets.

## Supporting Information Available

### Excel2SBOL

- GitHub: https://github.com/SynBioDex/Excel-to-SBOL
- Documentation: https://github.com/SynBioDex/Excel-to-SBOL/wiki
- Plugin: https://github.com/SynBioHub/Plugin-Submit-Excel2SBOL
- PyPI: https://pypi.org/project/excel2sbol/

### SBOL2Excel

- GitHub: https://github.com/SynBioDex/SBOL-to-Excel
- Documentation: https://github.com/SynBioDex/SBOL-to-Excel/wiki
- Plugin: https://github.com/SynBioHub/Plugin-Download-SBOL2Excel
- PyPI: https://pypi.org/project/sbol2excel/

### Suplemental Filesl

- Template for Excel to SBOL Conversion SBOL2 simple parts template.xlsx
- Output of SBOL to Excel sbol2excel output.xlsx
- Case Study Excel to SBOL (Experimental Data) flapjack compiler sbol3 v023.xlsx
- Case Study SBOL to Excel (Cello) cello.xlsx

